# Intraspecific variation in indirect plant-soil feedbacks as a driver of a wetland plant invasion

**DOI:** 10.1101/160523

**Authors:** Warwick J. Allen, Laura A. Meyerson, Andrew J. Flick, James T. Cronin

## Abstract

Plant-soil feedbacks (PSFs) can influence plant competition via direct interactions with pathogens and mutualists or indirectly via apparent competition/mutualisms (i.e., spillover to cooccurring plants) and soil legacy effects. Presently, it is unknown how intraspecific variation in PSFs interacts with the environment (e.g., nutrient availability) to influence competition between native and invasive plants. We conducted a fully crossed multi-factor greenhouse experiment to determine the effects of soil biota, interspecific competition, and nutrient availability on biomass of replicate populations from one native and two invasive lineages of common reed (*Phragmites australis*) and a single lineage of native smooth cordgrass (*Spartina alterniflora*). Harmful soil biota consistently dominated PSFs involving all three *P. australis* lineages, reducing biomass by 10%, regardless of nutrient availability or *S. alterniflora* presence as a competitor. Spillover of soil biota derived from the rhizosphere of the two invasive *P. australis* lineages reduced *S. alterniflora* biomass by 7%, whereas soil biota from the native *P. australis* lineage increased *S. alterniflora* biomass by 6%. Interestingly, regardless of lineage, *P. australis* soil biota negatively affected *S. alterniflora* biomass when grown alone (i.e., a soil legacy), but had a positive impact when grown with *P. australis*, suggesting that *P. australis* is preferred by harmful generalist soil biota or facilitates *S. alterniflora* via spillover (i.e., apparent mutualism). Soil biota also reduced the negative impacts of interspecific competition on *S. alterniflora* by 13%, although it remained competitively inferior to *P. australis* across all treatments. Moreover, competitive interactions and the response to nutrients did not differ among *P. australis* lineages, indicating that interspecific competition and nutrient deposition may not be key drivers of *P. australis* invasion in North America. Taken together, although soil biota, interspecific competition, and nutrient availability appear to have no direct impact on the success of invasive *P. australis* lineages in North America, indirect spillover and soil legacies from *P. australis* occur and may have important implications for co-occurring native species and restoration of invaded habitats. Our study integrates multiple factors linked to plant invasions, highlighting that indirect interactions are likely commonplace in driving successful invasions and their impacts on the local community.

## INTRODUCTION

Plant species influence the community composition and function of soil biota, which in turn can impact fitness of host plant species, a reciprocal interaction commonly referred to as a plant-soil feedback (PSF) (Kulmatiski et al. 2008). The net impact of PSFs on host plants depends on the balance between beneficial (nitrogen-fixing bacteria, mycorrhizal fungi, and other mutualists) and harmful (soil-borne pathogens, parasites, and herbivores) interactions with soil biota (Klironomos 2002, Reinhart and Callaway 2006). PSFs have clear implications for the success of invasive plant species (van der Putten et al. 2013). For example, invasive plants could experience less positive (i.e., weaker associations with mutualists) or more negative (i.e., greater attack by local natural enemies) PSFs relative to closely-related native species, suggesting some biotic resistance of the native soil community (Elton 1958, Callaway et al. 2013). Alternatively, invasive plants may generate more positive or less negative PSFs than closely-related native species, potentially resulting in dominance for the invader. Several empirical studies, metaanalyses, and reviews support this latter scenario (Klironomos 2002, Agrawal et al. 2005, Kulmatiski et al. 2008, Suding et al. 2013). Interestingly, generalist soil biota cultivated by invaders also interact with co-occurring native plant species, resulting in indirect effects of the invasive species mediated through PSFs (i.e., pathogen/mutualist spillover, more generally known as apparent competition/mutualism) (Eppinga et al. 2006, Mangla et al. 2008). Moreover, other plant species may also be inhibited by soil biota even after removal of the invasive plant (i.e., a soil legacy) (Corbin and D’Antonio 2012, Grove et al. 2015).

Little is known about how PSFs interact with other factors linked to plant invasions such as competitive interactions with native species and nutrient availability. For example, modeling and experimental studies have demonstrated that even low strength PSFs can alter interspecific competition (Bever 2003, Casper and Castelli 2007, Hodge and Fitter 2013), itself a key mechanism thought to underlie the success of invasive species (reviewed by Gioria and Osborne 2014). Likewise, anthropogenic nutrient deposition is a major component of global environmental change and a facilitating factor in many plant invasions (Dukes and Mooney 1999). Nutrient availability can alter competitive interactions (Wilson and Tilman 1993), activity of plant mutualists and pathogens in the soil (Johnson et al. 2008), and thus the direction and magnitude of PSFs (Manning et al. 2008). However, such a multi-factor approach has rarely been used to study the role of PSFs in invasion success (but see Larios and Suding 2015).

Finally, intraspecific genetic variation is an important part of ecological and evolutionary processes (see Bolnick et al. 2011 for review) and is known to alter the effects of nutrients (e.g., Saltonstall and Stevenson 2007) and competitors (e.g., Howard et al. 2008, Gomola et al. 2017) on plant fitness, and to influence community composition of soil biota (Schweitzer et al. 2008, Nelson and Karp 2013, Lamit et al. 2016, Bowen et al. in review), yet experiments examining intraspecific variation in PSFs remain rare (but see Bukowski and Petermann 2014, Maron et al. 2015, Wagg et al. 2015). Biological invasions are often characterized by multiple introduction events, which can lead to the presence of multiple genetic lineages in the introduced range (e.g., Durka et al. 2005, Meyerson et al. 2012, Gomola et al. 2017). Cryptic invasions have also been described (e.g., Saltonstall 2002, Tyler et al. 2007), where invasive genotypes or hybrids co-occur with native genotypes. Because many studies of invasive species assume that no intraspecific variation exists in their interactions with the environment and resident community, this can result in misleading insights and management recommendations, particularly if different genotypes employ different mechanisms of invasion (Meyerson and Cronin 2013, Gomola et al. 2017).

The goal of this study was to investigate the effects of plant intraspecific genetic variation, soil biota, and nutrient availability on competitive interactions between common reed, *Phragmites australis* (Cav.) Trin. ex Steudel, and a dominant, co-occurring, native marsh grass species, smooth cordgrass (*Spartina alterniflora* Loisel.). In North America, there is a widespread endemic native lineage of *P. australis* as well as two invasive lineages (European and Gulf) (Saltonstall 2002, Meyerson et al. 2009, Lambertini et al. 2012, Meyerson et al. 2012). We grew nine *P. australis* populations (three of each lineage) and a single population of *S. alterniflora* in pots containing live or sterilized soil inoculum from the rhizosphere of the respective *P. australis* population, at two nutrient levels, and with the two plant species either alone or together. Based on invasion biology dogma, we tested the following predictions: 1) invasive *P. australis* lineages have more positive PSFs than the native lineage; 2) indirect spillover and soil legacies of soil biota on *S. alterniflora* are more negative from invasive than native *P. australis* lineages; 3) the direction and strength of PSFs, spillover, and soil legacies depend upon the presence of an interspecific competitor and nutrient availability; 4) nutrient availability and lineage-specific PSFs alter interspecific competition between *P. australis* and *S. alterniflora*; and 5) invasive *P. australis* lineages exhibit stronger interspecific competitive ability and response to nutrient availability than native lineages and *S. alterniflora*.

## MATERIALS AND METHODS

### Study organisms

*Phragmites australis* is a model organism for studying plant invasions (Meyerson et al. 2016) and is one of the most widely distributed plants in the world. Multiple genetic lineages of *P. australis* grow sympatrically in North America (Saltonstall 2002, Meyerson et al. 2009, Lambertini et al. 2012, Meyerson et al. 2012, Meyerson and Cronin 2013). The native lineage is endemic to North America and consists of at least fourteen different haplotypes (Saltonstall 2002, Meadows and Saltonstall 2007). An invasive lineage of *P. australis* from Europe has spread aggressively in wetlands of North America since first appearing in herbarium records ~150 years ago (Saltonstall 2002, Howard et al. 2008, Meyerson et al. 2012). This invasive European lineage is mostly comprised of a single haplotype (*M*) and forms large, dense, monospecific populations that negatively impact hydrology, native plant diversity, habitat quality for fauna, and ecosystem function (reviewed by Meyerson et al. 2009). An additional lineage (known as Gulf) is widely distributed along the Gulf of Mexico and west to California (Lambertini et al. 2012, Meyerson et al. 2012) and is likely a recent arrival from Mexico or Central America (Colin and Eguiarte 2016). Although its mode of introduction to North America remains unknown, we classify it as invasive (following Richardson et al. 2000) due to its rapidly-growing populations (Bhattarai and Cronin 2014) and the speed with which it has spread (Meyerson et al. 2012).

Recent studies with *P. australis* have described distinct oomycete, archaea, and bacteria communities from rhizosphere soil of native and European *P. australis* lineages in North America (Nelson and Karp 2013, Crocker et al. 2015, Yarwood et al. 2016, Bowen et al. in review), suggesting that the net impact of soil biota (i.e., PSF) may also differ among *P. australis* lineages. However, virtually all studies to date have focused on describing community structure of soil biota, whereas the direction and magnitude of their impact on each *P. australis* lineage remain relatively unknown. The exception is the study by Crocker et al. (2015), which demonstrated that virulence of some isolated *Pythium* oomycetes differed between native and European lineages. Furthermore, the ecology, trophic interactions, or microbial community of the Gulf lineage remains virtually unknown (but see Bowen et al. in review).

The widely distributed perennial grass *S. alterniflora* is native to salt marshes on the East and Gulf Coasts of North America, but invasive in other locations, such as the West Coast of North America (Tyler et al. 2007) and China (Zhao et al. 2010, Li et al. 2014). We selected *S. alterniflora* as a standardized competitor because it is a dominant plant in many coastal marshes, where it co-occurs with *P. australis* (Bertness 1991, Medeiros et al. 2013) and even shares pathogens (Li et al. 2014). The response to abiotic factors and competitive ability of *S. alterniflora* have been well described; specifically, *S. alterniflora* has a strong positive response to increased nutrient availability (Tyler et al. 2007, Zhao et al. 2010) and is generally an inferior competitor to *P. australis* and other salt marsh plants, except in environments with high abiotic stress (Bertness 1991, Pennings et al. 2005, Medeiros et al. 2013).

### Greenhouse experiment design

We conducted a greenhouse experiment to examine the interactive effects of soil biota, interspecific competition, and nutrient availability on clonal growth (i.e., above- and belowground biomass) of the three *P. australis* lineages in North America and native *S. alterniflora*. The experimental design consisted of four treatments: 1) live/sterile soil biota - live or sterilized soil inoculum collected *in situ* from the rhizosphere of each *P. australis* population was added to each pot (10% of total soil weight to minimize variation in abiotic soil properties and nutrient flushes following soil sterilization). Soil biota was always combined with its associated *P. australis* population such that no mixing of soil and *P. australis* sources occurred. 2) presence/absence of an interspecific competitor - pots were planted with either *P. australis, S. alterniflora*, or both species combined. 3) high/low nutrient levels - nutrient levels were manipulated to represent nutrient-rich and nutrient-poor environments. 4) *P. australis* lineage - plants and corresponding soil inoculum from populations of the native, European, and Gulf lineages of *P. australis* were used for the experiment. These four treatments were fully crossed (36 total treatment combinations) and replicated among clones from three distinct *P. australis* populations within each lineage (Table 1). We planted ten replicates of each of the treatment combinations for all nine *P. australis* populations, resulting in a total of 1,080 pots. Plants were grown in a greenhouse located at Louisiana State University (30.36° N, −91.14° W) and pots were arranged in a randomized blocked design with five blocks to account for possible gradients in the greenhouse environment. A more detailed description of the experimental treatments and design is provided in Appendix S1.

**Table 1.** List of *Phragmites australis* field populations used for the greenhouse experiment.

| Population name, state (ID code) | Latitude | Longitude | Lineage | Status |
| --- | --- | --- | --- | --- |
| Palm Canyon Road, CA (PCN) | 33.83 | -116.62 | Native | Endemic |
| Little Caliente Hot Springs, CA (LCN) | 34.54 | -119.62 | Native | Endemic |
| Mackay Island, NC (NCN) | 36.51 | -75.95 | Native | Endemic |
| East Cameron, LA (ECM) | 29.77 | -93.29 | European | Invasive |
| I-40, AZ (I40M) | 34.72 | -114.49 | European | Invasive |
| Mackay Island, NC (NCM) | 36.51 | -75.95 | European | Invasive |
| Okeeheelee Park, FL (FLI) | 26.65 | -80.16 | Gulf | Invasive |
| Intracoastal City, LA (ICI) | 29.78 | -92.20 | Gulf | Invasive |
| Creole, LA (CRI) | 29.83 | -93.11 | Gulf | Invasive |

### Data collection

Harvesting was completed from 5 to 13 December, 2015. At this southern climate, plants were still growing and had not reached the flowering stage, which generally follows the second year of growth or later when propagating from rhizome cuttings. Above- and belowground biomass was harvested for each plant species from each pot, oven-dried to constant mass at 60 °C, and weighed to the nearest 0.1 g. Because no plants produced a panicle, we used above- and belowground biomass (i.e., clonal growth) and root:shoot ratio (i.e., biomass allocation) as proxies for fitness. As these variables all demonstrated similar results, we focus on total biomass for our results and discussion, but report fully on other variables in Appendices S3, S4, and S5.

### Data analysis

To examine how response variables for each plant species (*P. australis*, *S. alterniflora*) were influenced by *P. australis* lineage (native, European, Gulf), live/sterile soil inoculum, presence/absence of an interspecific competitor, and high/low nutrient availability, we used Akaike’s Information Criteria corrected for finite sample size (AICc) to select the most informative mixed-effect model from a set of candidate models (Burnham and Anderson 2010). The full model included the variables above and all two-, three-, and four-way interactions as fixed effects (fifteen total variables). *Phragmites australis* population and greenhouse block were included as random effects to account for within-lineage variation and possible greenhouse environment gradients, respectively. We report AICc weights that indicate the proportional strength of support for model *i* being the best model given the set of plausible models (ΔAICc ≤ 2). For our interpretations, we estimated least-squares means (back-transformed) based on the most likely model for each response variable and focused on effect sizes (i.e., proportional differences in means) (Burnham and Anderson 2010). For brevity, only results for models with AICc weight ≥ 0.30 are reported (i.e., the top model for each response variable) and we focus the discussion on the interesting yet poorly-understood interaction effects (see Appendix S2).

## RESULTS

### Factors affecting total biomass of Phragmites australis

Soil inoculum, interspecific competition, and nutrient availability were identified as influential in explaining variation in *P. australis* total biomass using AICc model selection. Four candidate models received adequate support (ΔAICc ≤ 2) and all included these same three main effects (cumulative AICc weight = 1) or various interactions between them. The top model (AICc weight = 0.429, Table 2) included the main effects only and had two times the support of the other three models (second top model: AICc weight = 0.218).

**Table 2.** AICc best models (ΔAICc ≤ 2) explaining variation in total biomass (square root transformed) for each plant species (*Phragmites australis* or *Spartina alterniflora*). Explanatory variables: L = *P. australis* lineage (native, European, Gulf), C = presence/absence of an interspecific competitor, N = high/low nutrient availability, and S = live/sterile soil inoculum. × denotes interactions between explanatory variables.

| Response variables | Models | AICc | $\Delta\text{AICc}$ | AICc weight |
| --- | --- | --- | --- | --- |
| *Phragmites australis* |  |  |  |  |
| Total biomass | C N S | 2611.6 | 0.00 | 0.429 |
| | C N S C×S | 2612.9 | 1.35 | 0.218 |
| | C N S N×S | 2613.3 | 1.73 | 0.181 |
| | C N S C×N | 2613.4 | 1.82 | 0.173 |
| *Spartina alterniflora* |  |  |  |  |
| Total biomass | C L N S C×N C×S L×N L×S | 2373.5 | 0.00 | 0.714 |
| | C L N S C×N C×S L×N L×S N×S C×N×S | 2375.4 | 1.83 | 0.286 |

Based on the top model, average biomass was 10% lower for *P. australis* grown in pots containing live (43.47 ± 0.5 g, back-transformed least-squares mean ± S.E.) than sterile soil inoculum (48.47 ± 0.5 g) (Fig. 1A), regardless of the *P. australis* lineage, presence of an interspecific competitor, or nutrient availability (no influential interactions in the top model). Similarly, competition with *S. alterniflora* reduced biomass of *P. australis* by 13% (42.72 ± 0.5 g) relative to when grown alone (49.27 ± 0.5 g) (Fig. 1B), but this effect did not differ among lineages or interact with soil biota or nutrient availability. Finally, although high nutrient availability doubled biomass production of *P. australis* (from 31.26 ± 0.5 g to 63.44 ± 0.5 g, Fig. 1C), this effect also did not interact with the other variables.

**Figure 1.**
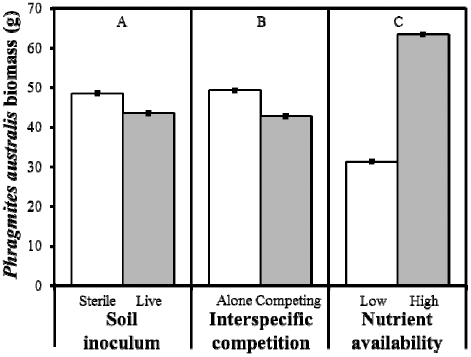
Influential main effects on *Phragmites australis* biomass (g) (least squares mean ± S.E.) identified using mixed-effect model selection: A) live or sterilized *P. australis* soil inoculum, B) presence or absence of competition with *Spartina alterniflora*, and C) high or low nutrient availability. No interactions among main effects were included in the top model (see Table 2).

### Factors affecting total biomass of Spartina alterniflora

The variables influential in explaining *S. alterniflora* biomass were: *P. australis* lineage, soil inoculum, interspecific competition, nutrient availability, and the lineage × soil inoculum, lineage × nutrient availability, interspecific competition × soil inoculum, and interspecific competition × nutrient availability interactions (top model: AICc weight = 0.714, Table 2). The second top model (AICc weight = 0.286) included these variables and additional interactions but had less than half the support of the top model.

Most interestingly, the direction of the impact of *P. australis* soil inoculum on *S. alterniflora* biomass depended upon the *P. australis* lineage the soil inoculum was sourced from (lineage × soil inoculum interaction) as well as the presence/absence of *P. australis* as a competitor (interspecific competition × soil inoculum interaction). The impact of soil biota on *S. alterniflora* biomass was in the opposite direction for invasive (a 7% decrease relative to sterile soil; European: 23.32 ± 0.06 g to 20.29 ± 0.06 g; Gulf: 23.78 ± 0.06 g to 20.68 ± 0.06 g) and native (a 6% increase from 18.14 ± 0.06 g to 20.44 ± 0.06 g) *P. australis* lineages, an overall difference in biomass of 13% (Fig. 2). Moreover, when grown alone, *S. alterniflora* biomass was 12% lower in pots with live (28.63 ± 0.04 g) than sterile (32.67 ± 0.04 g) *P. australis* soil inoculum (Fig. 3A). Conversely, when competing with *P. australis*, *S. alterniflora* plants in live soil inoculum had 6% higher biomass (13.68 ± 0.04 g) than those in sterile inoculum (12.92 ± 0.04 g), an 18% difference between treatments. Soil biota from *P. australis* also altered the effects of interspecific competition, reducing *S. alterniflora* biomass by 52% in live soil inoculum (from 28.63 ± 0.04 g to 13.68 ± 0.04 g) and 60% in sterile soil inoculum pots (from 32.67 ± 0.04 g to 12.92 ± 0.04 g) (Fig. 3A). The effects of interspecific competition also interacted with nutrient availability; biomass of *S. alterniflora* decreased by 53% (from 17.37 ± 0.04 g to 8.09 ± 0.04 g) and 58% (from 47.60 ± 0.04 g to 19.79 ± 0.04 g) in nutrient-poor and nutrient-rich pots, respectively (Fig. 3B). Increased nutrient availability had a strong effect on *S. alterniflora* biomass, which was 174% (from 17.37 ± 0.04 g to 47.60 ± 0.04 g) and 145% (from 8.09 ± 0.04 g to 19.79 ± 0.04 g) higher than in nutrient-poor pots when grown alone and with *P. australis* as a competitor, respectively (interspecific competition × nutrient availability interaction) (Fig. 3B). Finally, in nutrient-poor pots, differences in *S. alterniflora* biomass among pots with soil inoculum from different *P. australis* lineages were relatively small (< 4%, range of 12.10 to 12.52 ± 0.06 g). However, in nutrient-rich pots, *S. alterniflora* grown with soil inoculum from the invasive lineages of *P. australis* had 19% higher biomass (European: 34.30 ± 0.06 g; Gulf: 34.62 ± 0.06 g) than pots with soil inoculum or plants from the native lineage (27.89 ± 0.06 g) (lineage × nutrient availability interaction). This pattern was consistent regardless of the presence of *P. australis* as a competitor or whether the soil inoculum was live or sterile.

**Figure 2.**
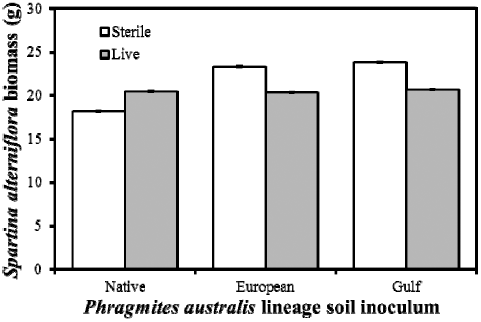
Indirect impact of live or sterilized soil inoculum obtained from the three *Phragmites australis* lineages on biomass (g) (least squares mean ± S.E.) of *Spartina alterniflora*. The interaction between soil inoculum and *P. australis* lineage was identified as influential using mixed-effect model selection (see Table 2). Some error bars are obscured due to their small size.

**Figure 3.**
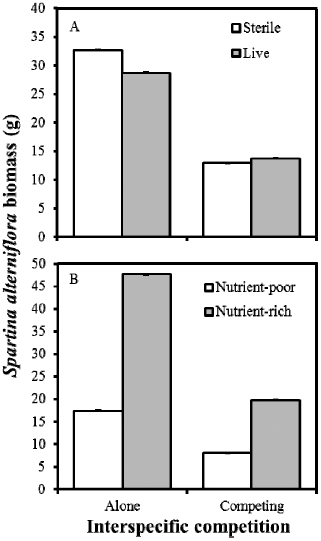
Influential interaction effects of competition with *Phragmites australis* on biomass (g) (least squares mean ± S.E.) of *Spartina alterniflora* grown in A) live or sterilized *P. australis* soil inoculum and B) with high or low nutrient availability. These interactions were identified as influential using mixed-effect model selection (see Table 2). Error bars are obscured due to their small size.

### Factors affecting other response variables

The top models for other *P. australis* and *S. alterniflora* response variables (aboveground biomass, belowground biomass, root:shoot ratio, see Appendices S3, S4, and S5) were remarkably similar to those for total biomass. However, lineage-specific effects were more prevalent and three additional terms were identified as influential, which we focus on describing below. For *P. australis*, the negative impact of live soil biota on aboveground biomass of the European lineage was 64% and 75% less than the native and Gulf lineages (lineage × soil inoculum interaction), respectively (Appendix S5, Fig. S1). The European lineage also had the greatest root:shoot ratio in nutrient-poor pots, but this changed to the native lineage in nutrient-rich pots (lineage × nutrient availability interaction) (Appendix S5, Fig. S2). Finally, for the additional *S. alterniflora* traits, the tripartite lineage × interspecific competition × soil inoculum interaction was the only other variable identified as influential that was not already present in the top model for total biomass. For this interaction, the presence of *P. australis* increased the strength of soil biota impacts on *S*. *alterniflora* root:shoot ratio across all three lineages, but the direction of this effect varied by *P. australis* lineage (Gulf: a 3% to 34% decrease in root:shoot ratio; European: 6% to 23% decrease; native: 9% to 27% increase) (Appendix S5, Fig. S3).

## DISCUSSION

Our study suggests that interactions with soil biota do not directly influence the success of invasive *P. australis* lineages, with this view supported by a similar reduction (10%) in total biomass of native and invasive (European and Gulf) lineages through interactions with their respective soil biota, irrespective of the presence of an interspecific competitor or nutrient availability. `This consistent negative impact of soil biota on *P. australis* supports the established view that conspecific PSFs are predominantly negative (Bever 2003, Kulmatiski et al. 2008). On the other hand, soil biota from invasive *P. australis* populations reduced biomass of native *S. alterniflora*, whereas soil biota from native *P. australis* populations had a positive effect, suggesting the potential to exclude and facilitate co-occurring native plant species, respectively (Bever et al. 1997, Klironomos 2002, van der Putten et al. 2013). Interestingly, PSFs involving *P. australis* soil biota were negative for *S. alterniflora* grown alone (i.e., a negative soil legacy) but positive when grown in the presence of *P. australis*, suggesting that *P. australis* may be preferred by harmful generalist soil biota, or facilitates *S. alterniflora* via apparent mutualism. To our knowledge, this is the first study to demonstrate that the direction of indirect PSFs differs among conspecific native and invasive plant taxa, and can change depending upon the presence/absence of the initial host plant of the soil biota (i.e., between spillover and soil legacy). Consistent with previous studies (Bertness 1991, Pennings et al. 2005), we also found *P. australis* to be a dominant interspecific competitor and that *S. alterniflora* had a stronger response to increased nutrient availability, which also mediated the impact of interspecific competition on *S. alterniflora*. However, contrary to expectations, we found little support for the hypothesis that invasive *P. australis* lineages have superior interspecific competitive ability or response to nutrients when compared to the native lineage. Taken together, these results suggest that rather than direct PSFs, interspecific competitive ability, and response to nutrient deposition, the more nuanced indirect effects of *P. australis* soil biota on *S. alterniflora* via spillover of pathogens and mutualists (i.e., apparent competition and mutualism), soil legacy effects, and altered interspecific competition strength appear to be the key differences among *P. australis* lineages that may ultimately influence their relative success in North American salt marshes. Finally, our study, which integrates multiple factors linked to invasion success, highlights how indirect interactions can underpin successful invasions and their impact, and could inform approaches to management and restoration of areas invaded by *P. australis*.

### Direct PSFs of Phragmites australis lack intraspecific variation or context dependency

Despite strong lineage-specific differences in rhizosphere microbial communities (Nelson and Karp 2013, Yarwood et al. 2016, Bowen et al. in review), impact of *Pythium* spp. pathogens (Crocker et al. 2015), and soil biota effects on aboveground biomass (this study), we found that the impact of soil biota was consistently negative for all three *P. australis* lineages - a 10% reduction in total biomass. Thus, contrary to previous studies (Klironomos 2002, Agrawal et al. 2005, Kulmatiski et al. 2008, Suding et al. 2013, but see Callaway et al. 2013) and our first prediction, invasive *P. australis* lineages do not benefit from more positive PSFs than the native lineage, indicating that soil biota does not directly facilitate the relative success of invasive *P. australis* lineages in North America. This unexpected result is consistent with that of Bowen et al. (in review), who used structural equation modeling to show that rhizosphere bacterial richness, activity, and metabolism did not mediate *P. australis* biomass. One possible reason for the lack of differences in PSF strength among lineages could simply be that although lineages differ in their microbial communities, their net effects on the plant are the same. However, studies in other systems contradict this explanation, such as that of Wagg et al. (2015) who demonstrated that differences in PSFs of two populations of *Trifolium pratense* were related to corresponding differences in the rhizosphere microbe community. Finally, in contrast to our third prediction, the impact of soil biota on *P. australis* biomass was unaffected by the presence of *S. alterniflora* as a competitor or nutrient availability, suggesting there is little context dependency of *P. australis* PSFs regarding these variables. Intriguingly, these results are similar to those of the only other study to take such a multi-factor approach to the role of PSFs in plant invasions, where Larios and Suding (2015) found that negative PSFs of invasive wild oat (*Avena fatua*) were also largely unaffected by competition or nutrient availability.

### Lineage-specific indirect PSFs influence Phragmites australis invasion and impact

In support of our second prediction, we found that generalist soil biota from the rhizosphere of populations of the two invasive *P. australis* lineages had a net negative impact on *S. alterniflora* biomass, whereas soil biota from populations of the native lineage had a net positive impact, regardless of the presence of *P. australis* plants or nutrient availability. The large extent and density of populations of the invasive *P. australis* lineages relative to the native lineage and other native wetland plants means that even small invasion-induced changes in PSFs could be widespread and important in invaded habitats. Thus, our study represents the first to demonstrate intraspecific variation in spillover and provides support for its importance as a potential mechanism driving plant invasions. One possible explanation for the negative impact on *S. alterniflora* could be that invasive *P. australis* accumulates local generalist soil pathogens, which spillover onto *S. alterniflora*, overwhelming any positive impacts from beneficial organisms (Borer et al. 2007, Mangla et al. 2008). Similarly, beneficial soil biota may spillover to *S. alterniflora* from soil associated with the native *P. australis* lineage, representing a possible explanation for why the native *P. australis* lineage usually co-occurs with a diverse suite of other native species (Meyerson et al. 2009). These indirect interactions are representative of apparent competition and mutualism, respectively, whereby shared natural enemies or mutualists mediate interactions between two or more species. There is growing support for apparent competition involving herbivores and pathogens as an important driver of plant invasions (Borer et al. 2007, Dangremond et al. 2010), including for *P. australis* (Bhattarai et al. 2017a). Interestingly, Li et al. (2014) demonstrated pathogen spillover between *P. australis* and *S. alterniflora* in the Yangtze River estuary in China, but the roles of the species were reversed; there, *S. alterniflora* is invasive and spillover of the fungal pathogen *Fusarium palustre* resulted in significant dieback of native *P. australis*. Unfortunately, our experimental design did not allow us to assess how soil biota sourced from *S. alterniflora* populations affects *P. australis* in North America, although a reciprocal transplant experiment using soil biota from *S. alterniflora* and other native plants is a logical next step.

Furthermore, live *P. australis* soil biota decreased native *S. alterniflora* biomass by 12% when grown alone, but this changed to a 6% increase when competing with *P. australis*. We observed similar interspecific competition × soil inoculum interaction effects for all *S. alterniflora* response variables, although the direction of this impact on biomass allocation (root:shoot ratio) also differed among lineages, further supporting our predictions of strong interplay among these three factors. Similarly, Larios and Suding (2015) demonstrated that native purple needlegrass (*Stipa pulchra*) exhibited a positive PSF when grown alone at low nutrient levels, but that this PSF became neutral when competing with *A. fatua* or in high nutrient environments. At least two scenarios could explain the effects we observed for *S. alterniflora*. First, harmful generalist soil biota may interact preferentially with *P. australis*, only switching hosts to *S. alterniflora* when *P. australis* is absent. Such a preference is not entirely unexpected given that the soil inoculum was originally collected from naturally occurring *P. australis* populations and likely contained organisms coadapted to that population, lineage, and species (Nelson and Karp 2013, Yarwood et al. 2016, Bowen et al. in review). Thus, we suggest that *P. australis* generates a negative soil legacy whereby harmful generalist soil biota switch to native host species when *P. australis* is no longer present. Negative soil legacies appear to be relatively common among invasive species and are widely-recognized to prevent establishment of native plants and improve chances of invader recolonization (D’Antonio and Meyerson 2002, Corbin and D’Antonio 2012, Grove et al. 2015). Second, our findings could be indicative of spillover of beneficial soil biota cultivated by *P. australis* to *S. alterniflora* (i.e., an apparent mutualism), suggesting that *P. australis* may indirectly facilitate the growth of co-occurring native plants. However, these underlying mechanisms cannot easily be disentangled without identifying the organisms involved, which was unfortunately outside the scope of this study.

### Effects of interspecific competition and nutrient availability

Superior competitive ability has long been recognized as a common trait of invasive plant species (Elton 1958, reviewed by Gioria and Osborne 2014) and is often cited as one of the main reasons the European *P. australis* lineage has become so prevalent in North America. In support of this view and our fifth prediction, we found that interspecific competition decreased biomass of *P. australis* and *S. alterniflora* by 13% and 57%, respectively. This result is consistent with studies showing that native *S. alterniflora* tends to be restricted to lower marsh areas due to its poor competitive ability but superior tolerance of abiotic stress factors such as high salinity and flooding (Bertness 1991, Pennings et al. 2005). Several studies have also indicated that European *P. australis* is a stronger competitor than the native and Gulf lineages (Howard et al. 2008, Holdredge et al. 2010). However, we failed to find any differences in total biomass, interspecific competitive ability, or impact on *S. alterniflora* biomass among the three *P. australis* lineages. Thus, we suggest that interspecific competitive ability may not be a key factor explaining the predominance of European relative to native and Gulf *P. australis* in North America.

Several studies have found that soil biota and nutrient availability can significantly alter the outcome of interspecific competition (Casper and Jackson 1997, Casper and Castelli 2007, Hodge and Fitter 2013, but see Maron et al. 2016). In our study, live soil biota and nutrient-poor conditions both reduced the negative impact of interspecific competition on biomass of *S. alterniflora* but not *P. australis*, partially supporting our fourth prediction. The effect of soil biota on interspecific competition can likely be attributed to the consistent negative PSF suffered by *P. australis,* which may decrease its competitive ability or the strength of beneficial spillover affecting *S. alterniflora*. Moreover, our findings contrast with earlier studies that found nutrient addition reduces negative impacts of interspecific competition on *S. alterniflora* (Levine et al. 1998, Emery et al. 2001). However, these experiments did not use *P. australis* as a competitor, a species possessing one of the highest nitrogen use efficiencies of all land plants (Mozdzer et al. 2013). Furthermore, at high levels of nutrient availability, light becomes the main limiting resource in plant competition (Casper and Jackson 1997, Aerts 1999), meaning that the taller *P. australis* would continue to outcompete the shorter *S. alterniflora*. Perhaps most importantly, no differences in the impact of these competitive interactions on *S. alterniflora* total biomass were detected among *P. australis* lineages, indicating they are unlikely to be important in explaining the relative success of invasive versus native *P. australis* lineages in North America.

Increased nutrient deposition via disturbance and anthropogenic modification is considered a major contributing factor to *P. australis* invasion success (Bertness et al. 2002, Holdredge et al. 2010) and plant invasions in general (Dukes and Mooney 1999). Unsurprisingly, nutrient availability had a strong effect on biomass of both our study species, but this was greater for *S. alterniflora* than *P. australis* (Zhao et al. 2010), which may help explain why *S. alterniflora* has become an invasive plant in salt marshes on the West Coast of North America (Tyler et al. 2007), China (Zhao et al. 2010, Li et al. 2014), and elsewhere. However, the strength of nutrient effects did not vary among *P. australis* lineages, which could be considered unusual, given that European invasive *P. australis* enjoys a higher maximum nutrient uptake ability than the native lineage (Mozdzer et al. 2010). However, differences may be more subtle, such as the stronger plasticity in biomass allocation (root:shoot ratio) in response to nutrient addition that we observed for the European invasive lineage, which may impact other measures of fitness (i.e., sexual reproduction) or biomass over more than one growing season. Additionally, *S. alterniflora* grown in pots containing soil inoculum from the native *P. australis* lineage did not respond as positively to nutrient additions as plants associated with soil inoculum from the invasive *P. australis* lineages. This effect was independent of the presence of *P. australis* and soil biota sterilization, suggesting that abiotic factors of the original soil inoculum may have affected nutrient uptake of *S. alterniflora* in nutrient-rich pots, which is surprising given the low soil inoculum ratio of 10% of total soil weight. Furthermore, the influence of nutrient availability on *P. australis* was unaffected by the presence of live soil biota and interspecific competitors, suggesting that the harmful effects of negative PSFs and interspecific competition do not impact the response of *P. australis* to changes in nutrient availability. For *S. alterniflora*, however, the biomass increase in response to nutrients was highest when grown on its own. This is unsurprising, given that *P. australis* is the superior competitor and its presence should reduce the ability of *S. alterniflora* to fully utilize resources. Importantly, this effect also did not vary among *P. australis* lineages, and taken together, our results suggest that nutrient deposition may not directly contribute to the spread of invasive *P. australis* lineages into wetlands and marshes dominated by native *P. australis* or *S. alterniflora*.

### Management implications and future directions

The identity and impact of the soil community should be an important consideration when attempting to restore habitat occupied by invasive plant species (D’Antonio and Meyerson 2002, Corbin and D’Antonio 2012). Thus, we suggest microbial inoculation (e.g., Middleton and Bever 2012), topsoil removal (e.g., Hölzel and Otte 2003), and planting of the native *P. australis* lineage or other native plants as potentially useful approaches to reduce the effects of harmful soil biota, promote cultivation of beneficial soil biota, and facilitate the development of a diverse native community in areas where invasive *P. australis* is being managed. Successful restoration may be crucial to preventing re-establishment of invasive *P. australis* by improving resistance to colonization by seedlings and vegetative spread (Byun et al. 2013). Management of the surrounding matrix to minimize propagule pressure from neighboring invasive *P. australis* populations would also be beneficial when considering such an approach. Furthermore, future studies should focus on the identification of lineage-specific pathogens and mutualists which may be useful in novel management efforts with the goal of controlling invasive *P. australis* lineages and restoring the native lineage, respectively (Kowalski et al. 2015). Finally, because invasive species interact directly and indirectly with a complex community of organisms and abiotic conditions, expanding PSF studies to multitrophic and community-level interactions, and continuing to address context dependency, is critical to furthering our understanding of the role of PSFs in plant invasions.

## ACKNOWLEDGEMENTS

We thank the organizations that granted permission to collect bulk soil: Mackay Island National Wildlife Refuge, Palm Beach County Parks Department, Los Padres National Forest, and Thousand Palms Oasis Preserve. Thanks also to G. Bhattarai, A. Lambert, C. Oster, R. Harman, N. Harms, J. Croy, M. Faldyn, M. Guedry, B. Becnel, E. Haydel, L. Nguyen, T. Tran, R. Daigle, K. Manley, M. McDaniels, M. McDonald, S. Campbell, M. Smith, and R. Mason for field, lab, and greenhouse assistance. This project was supported by NSF grants DEB 1050084 and DMS 1516833 (to J.T.C.), DEB 1501775 (to J.T.C. and W.J.A.), and DEB 1049914 (to L.A.M.). Additional support was provided to W.J.A. from the Louisiana Environmental Education Commission, Sigma Xi Scientific Research Society, and LSU Biograds.

